# Comparative analysis of corrected tiger genome provides clues to their neuronal evolution

**DOI:** 10.1101/544809

**Authors:** Parul Mittal, Shubham Jaiswal, Nagarjun Vijay, Rituja Saxena, Vineet K. Sharma

**Author notes:** Email address of authors: Parul Mittal –; Shubham K. Jaiswal –; Nagarjun Vijay –; Rituja Saxena –; Vineet K. Sharma –.

## Abstract

The availability of completed and draft genome assemblies of tiger, leopard, and other felids provides an opportunity to gain comparative insights on their unique evolutionary adaptations. However, genome-wide comparative analyses are very sensitive to errors in genome sequences and thus require accurate genomic assemblies for reliable evolutionary insights. In this study, while analyzing the tiger genome, we found almost one million erroneous substitutions in the coding and non-coding region of the genome affecting 4,472 genes, hence, biasing the current understanding of tiger evolution. Moreover, these errors produced several misleading observations in previous studies. Thus, to gain insights into the tiger evolution, we corrected the erroneous bases in the genome assembly and gene set of tiger, which was also validated by resequencing of a Bengal tiger genome and transcriptome. A comprehensive evolutionary analysis was performed using 10,920 orthologs from nine mammalian species including the corrected gene sets of tiger and leopard, and using five different methods at three hierarchical levels i.e. felids, Panthera, and tiger. The unique genetic changes in tiger revealed that the genes showing the signatures of adaptation in tiger were enriched in development and neuronal functioning. Specifically, the genes belonging to Notch signalling pathway, which is among the most conserved pathways involved in embryonic and neuronal development, were found to be significantly diverged in tiger in comparison to the other mammals. Our findings suggest the role of adaptive evolution in neuronal functions and development processes, which correlates well with the presence of exceptional traits such as sensory perception, strong neuro-muscular coordination, and hypercarnivorous behavior in tiger.

## INTRODUCTION

The advancement in genomic sequencing technologies has provided a tremendous impetus for studying the molecular and genetic basis of adaptive evolution. A recent accomplishment is the genome sequencing of tiger, the largest felid and a model species to identify the molecular adaptations to hypercarnivory ^1–3^. Tiger is a prominent member of the big cats, which are the topmost predators in the food chain, and play a key role in the ecological niche ^4^. It is a solitary animal with extraordinary muscle strength and predatory capabilities ^2,3^. The tiger genome sequencing revealed several molecular signatures of selection, particularly the rapid evolution in genes related to muscle strength, energy metabolism, and sensory nerves ^1^. Similar studies in felids, including the tiger, had also shown strong positive selection in genes related to sensory perception and neurotransmitters ^5^.

While genome sequences are indispensable for comparative genome-wide evolutionary studies, quality of a genome is crucial for such analyses and in deriving reliable inferences ^6–9^. The quality of a genomic assembly is commonly assessed based on the N50 values of contigs and scaffolds and does not account for single nucleotide errors, which are mainly introduced by the read error correction tools or *de novo* assembler ^10–15^. Such sequence errors in genomes can produce drastically misleading results in comparative genomic and evolutionary studies ^7,8^. We found a similar case in the tiger genome assembly reported by Cho et al. in 2013 ^1^ and available at Ensembl release 94 (PanTig1.0) ^16^ and NCBI. The presence of several erroneous single nucleotide substitutions in the assembly bias the current understanding of the tiger evolution.

Therefore, to perform a comprehensive genome-wide analysis of tiger, we sequenced the genome and transcriptome of a male Bengal tiger and corrected the errors in the earlier-reported tiger genome assembly. Using the corrected genome assembly and gene set of tiger, we carried out a comparative genomic analysis of tiger with several other mammalian species, which provided novel insights into the adaptive evolution of the lineage leading to tiger.

## METHODS

### Sample collection, DNA isolation, and sequencing of the Bengal tiger genome

Approximately 5-6 ml blood was drawn from the tail vein of a four years old male tiger at Van Vihar National Park, Bhopal, India and was collected in EDTA-coated vials. The fresh blood sample was immediately brought to the laboratory at 4 °C and genomic DNA was extracted using DNeasy Blood and Tissue Kit (Qiagen, USA) following the manufacturer’s protocol. Multiple shotgun genomic libraries were prepared using Illumina TruSeq DNA PCR-free library preparation kit and Nextera XT sample preparation kit (Illumina Inc., USA) as per the manufacturer’s instructions. The insert size for the TruSeq libraries was 350 and 550 bp, and the average insert size for Nextera XT libraries was ~650 bp. The insert size for both the libraries was assessed on 2100 Bioanalyzer using High Sensitivity DNA kit (Agilent, USA). The libraries were quantified using KAPA SYBR FAST qPCR Master mix with Illumina standards and primer premix (KAPA Biosystems, USA), and Qubit dsDNA HS kit on a Qubit 2.0 fluorometer (Life Technologies, USA) as per the recommended Illumina protocol. The normalised TruSeq 550 bp and Nextera XT libraries were loaded on Illumina NextSeq 500 platform using NextSeq 500/550 v2 sequencing reagent kit (Illumina Inc., USA) and 150 bp paired-end sequencing was performed. The TruSeq libraries of 350 bp were sequenced on Illumina HiSeq platform to generate 250 bp paired-end reads.

### RNA isolation and transcriptome sequencing

Total RNA extraction was carried out from the blood sample for transcriptomic analysis. The blood sample (~5 ml) was transferred into a 50 ml polypropylene conical centrifuge tube. The volume was brought up to 45 ml with 1x RBC Lysis Buffer (10x RBC Lysis Buffer: 89.9 g NH4Cl, 10.0 g KHCO3 and 2.0 ml 0.5 M EDTA dissolved in approximately 800 ml ddH2O and pH adjusted to 7.3) and incubated at room temperature for 10 minutes. The cells were pelleted at 600xg (~1,400 rpm) for 10 minutes in a room temperature centrifuge and the supernatant was discarded. The pellet was gently resuspended in 1 ml of RBC Lysis Buffer and transferred to a 1.5 ml microcentrifuge tube and incubated at room temperature for 5 minutes. The cells were pelleted for 2 minutes by centrifuging at room temperature at 3000 rpm. The supernatant was discarded, and the pellet was resuspended in 1 ml of sterile DPBS. The cells were again pelleted at room temperature at 3,000 rpm, and the supernatant was discarded. 1200 µl of TRIzol solution was added to each tube. 0.2 ml of chloroform was added, and the tube was vortexed for 15 seconds. The sample was then centrifuged at 13,000 rpm for 10 minutes at 4°C. The upper phase was removed and transferred to a clean microcentrifuge tube. To the remaining upper phase, an equal volume of cold isopropanol was added, and inverted to mix. The sample was placed in a −20°C freezer to precipitate. Sample was then centrifuged at 13,000 rpm for 10 minutes at 4°C. The supernatant was carefully discarded, and the pellet was rinsed with 0.5 ml of ice-cold 75% ethanol. The sample was centrifuged at 13,000 rpm for 10 minutes at 4°C. The supernatant was discarded, and the pellet was allowed to dry for 5 to 10 minutes to remove any remaining ethanol. The RNA pellet was dissolved by adding 20 µl of RNAse-free water. The transcriptomic libraries were prepared from the total RNA using the SMARTer universal low input RNA kit and TruSeq RNA sample prep kit v2 using the manufacturer’s instructions, and 100 bp paired end sequencing was performed on the Illumina HiSeq platform.

### Data download and preparation

For assembly correction, the latest assemblies of tiger (*Panthera tigris altaica*) and leopard (*Panthera pardus*) genome were retrieved from Ensembl release 94 (PanTig1.0 and PanPar1.0) ^16^. The reads data was retrieved from NCBI SRA with Accession SRX272981, SRX272988, SRX272991, SRX272997, SRX273000, SRX273020 and SRX273023 for Amur tiger. For leopard, the reads data with SRA Accession SRX1495683, SRX1495735, and SRX1495737 were retrieved from NCBI SRA. To construct the corrected CDS, the reference gtf was downloaded from Ensembl release 94 (Panthera pardus 1.0.93 and Panthera tigris altaica 1.0.93) ^16^. The retrieved raw reads of Amur tiger were mapped to the tiger reference assembly and raw reads of leopard were mapped to the leopard genome assembly using bwa mem (v0.7.12) ^17^ using default parameters. The alignment file was sorted and split scaffold-wise using Samtools (v1.4) ^18^.

### Genomic and CDS correction

The per-nucleotide metrics for each scaffold was calculated using bam-readcount tool (github/genome/bam-readcount) using minimum mapping quality 25, minimum base quality 25, and maximum depth 400. The base positions with less than 10x coverage or more than 200x coverage, or percent indel> 10% were filtered out to remove the low coverage, potentially repetitive, and indel regions, respectively, and the remaining bases were analyzed further. At a given position, if the representation of the reference base was less than one-fifth of the most frequent base, then the reference base was replaced by the most frequent base at that position. The adoption of this stringent criteria ensured that only those positions were corrected where a sufficient coverage of the most frequent base relative to the reference base was available to justify the replacement of the reference base. The above criteria were optimized after several iterations and visualization of randomly selected regions with the mapped reads in the IGV software ^19^. The corrected genome assembly was used to construct the corrected gene set using the gene structure information available at Ensembl.

### Orthologous gene set construction

An orthologous gene set was constructed using nine species – *Homo sapiens* (human), *Mus musculus* (mouse), *Bos taurus* (cat), *Equus caballus* (horse), *Canis familiaris* (dog), *Mustela putorius* (ferret), *Felis catus* (cat), *Panthera tigris altaica* (tiger) and *Panthera pardus* (leopard). The gene sets were retrieved from Ensembl release 94 (Cat: Felis_catus_9.0, Cow: UMD3.1, Dog: CanFam3.1, Horse: EquCab2, Human: GRCh38, Ferret: MusPutFur1.0, Mouse: Mus_musculus.GRCm38) ^16^. The corrected gene sets of tiger and leopard were used in the analysis. Information on one-to-one orthologs for the above species was retrieved from BioMart (Ensembl browser 94) ^20^.

#### Protein and nucleotide alignment

The one-to-one orthologs were filtered for the presence of premature stop codons (non-sense mutations). The gene phylogeny of each ortholog was inferred from the species phylogeny and was subjected to protein alignment using SATé-II ^21^, which implemented PRANK for alignment, Muscle for merging the alignment, and RAxML for tree estimation. The protein-based nucleotide alignment was carried out using ‘tranalign’ tool in EMBOSS package ^22^.

### Evolutionary analysis

#### Higher branch dN/dS

The variation in ω ratio between lineages on individual genes was calculated using the branch model in CodeML from the PAML software package (v4.9a) ^23^. The codons with any ambiguity site were removed from the analyses. The genes that qualified likelihood ratio test using a conservative 5% false-discovery-rate criterion against the null model (One ratio) were considered for further analysis. Also, the genes with dN/dS values >3 were not used for further analysis ^24,25^. The genes having a higher branch dN/dS values for foreground lineage compared to the background lineage were considered to show divergence (HBW: higher branch omega).

#### Positively selected genes and sites

To identify positively selected genes, a branch-site model was used in PAML software package (v4.9a) ^23^. The codons with any ambiguity site among the nine species were removed from the analyses. The genes that qualified the likelihood ratio test against the null model (fixed omega) with 5% false-discovery-rate were considered as positively selected genes (PSG). The sites with greater than 0.95 probability value for foreground lineages in Bayes Empirical Bayes analysis were considered as positively selected sites (PSS).

#### Unique substitutions and functional impact

Unique substitutions in amino acid in tiger were identified using the aligned protein sequences. The positions identical in all species but different in tiger were considered as a unique substitution in tiger. Any gap or unknown position was ignored. Five sites around any gap in the protein alignment were also ignored from the analysis. Unique substitutions in Panthera and felids were identified using the same approach. Functional impact of the substitutions were identified using Sorting Intolerant From Tolerant (SIFT) ^26^ tool and the UniProt database ^27^ was used for reference.

#### Higher nucleotide divergence

The maximum likelihood phylogenetic tree for each gene using its CDS alignments was constructed using PhyML package v3.1 ^28^. The root-to-tip branch length distances were calculated for each species using the ‘adephylo’ package in R ^29,30^. The genes with a significantly higher root-to-tip branch length for lineage leading to tiger compared to all other lineages were considered to show higher nucleotide divergence in tiger.

#### Identification of genes with multiple signs of adaptation

Genes showing more than two signs of adaptive divergence among the five signs (Unique substitution, higher dN/dS, positive selection, positively selected sites, and higher nucleotide divergence) used in the study were considered to be the genes with multiple signs of adaptation (MSA). Enrichment of MSA genes was carried out using WebGeStalt web server ^31^. The GO enrichment with p-value < 0.05 in over-representation enrichment analysis were considered to be enriched. The eggNOG analysis of the MSA genes was performed using the eggNOG v4.5.1 ^32^. The network-based pathway enrichment analysis was carried out based on the methodology implemented in EnrichNet ^33^ to identify the network interconnectivity score (XD-score) and classical overlap-based enrichment score (Fisher’s exact test adj. using Benjamini-Hochberg) using KEGG as the reference database. The significance threshold was calculated by performing a linear regression of network interconnectivity score (XD-score) and enrichment score (Fisher’s q-value). The pathways above the significance threshold were considered as enriched.

## RESULTS

Comparative genomic analysis was performed to gain insights into the evolution of tiger with several other mammalian species. Tiger is a prominent member of the Panthera genus, which is a fast evolving group that has undergone recent radiation with rapid functional diversification ^34–36^. Thus, the comparative genome-wide study of tiger with respect to the closely related Panthera species and other mammals is likely to provide novel evolutionary insights into their adaptive evolution. The tiger genome assembly, reported by Cho et al. in 2013 (available at http://tigergenome.org), was retrieved and used for the comparative analysis. During the analysis, we observed that the assembly comprised of several erroneous single base substitutions, which were perhaps introduced by the *de novo* assembler or by the read correction tools (**Supplementary Text S1**) ^10–15^. Similar errors were also present in the genome assembly of tiger, NCBI (GCA_000464555.1) and Ensembl (PanTig1.0). As observed from the analysis performed using the publicly available tiger genome assembly presented in the **Supplementary Text S1**, the above errors produced several misleading results (**Supplementary Figure S1-S2**). For example, BEX3, a gene that plays an important role in the neuronal apoptosis ^37,38^ was found to be positively selected and showed nine unique amino acid substitutions in tiger. Our analysis revealed that eight of the nine substitutions in this gene were due to the single nucleotide errors in the publicly available genome assembly of tiger, which produced the incorrect result shown in **Supplementary Figure S2.** We found several such cases in previous studies where the erroneous single nucleotide substitutions have led to incorrect evolutionary interpretations **(Supplementary Text S2, Figure S3-S4)**.

Therefore, to understand the adaptive evolution of tiger lineage, we first corrected the available reference assembly of the tiger genome, which was further validated by sequencing the genome and transcriptome of a Bengal tiger from India. A total of 175,680,850 paired-end reads (67 GB) and 32,252,904 reads (6 GB) were generated for the genome and transcriptome, respectively **(Supplementary Table S1-S2).** Among the five extant species in the Panthera genus, the genome assemblies are publicly available for only two species, tiger and leopard ^1,39^. Thus, in addition to tiger, we also generated the corrected genome assembly of leopard using the strategy shown in **Figure 1a**, and briefly mentioned below.

### Correcting the genome assembly and gene set of tiger and leopard

The genomic reads were mapped to the tiger genome assembly obtained from Ensembl (PanTig1.0), and the incorrect positions in the assembly were identified using the read alignments. We developed a pipeline, named ‘SeqBug’, for genome assembly corrections for single nucleotide errors introduced primarily by the read error correction or *de novo* assembler. The method is based on the mapping of high-depth short reads to the genome assembly followed by the identification of an incorrect base-pair and its correction. A given base-pair was considered ‘incorrect’ if it had 20% or less representation than any other base in the read alignments among all the mapped reads. The detailed criteria used for base correction is provided in **Methods**. A total of 982,606 bases (0.04% of the genome) were corrected in tiger assembly. The distribution of per-base coverage of these corrected positions showed a Poisson distribution (with a peak on the low coverage side) in comparison to the normal distribution for the whole genome (**Figure 1b**), suggesting that a larger number of corrections were made in the low coverage regions.

**Figure 1.**
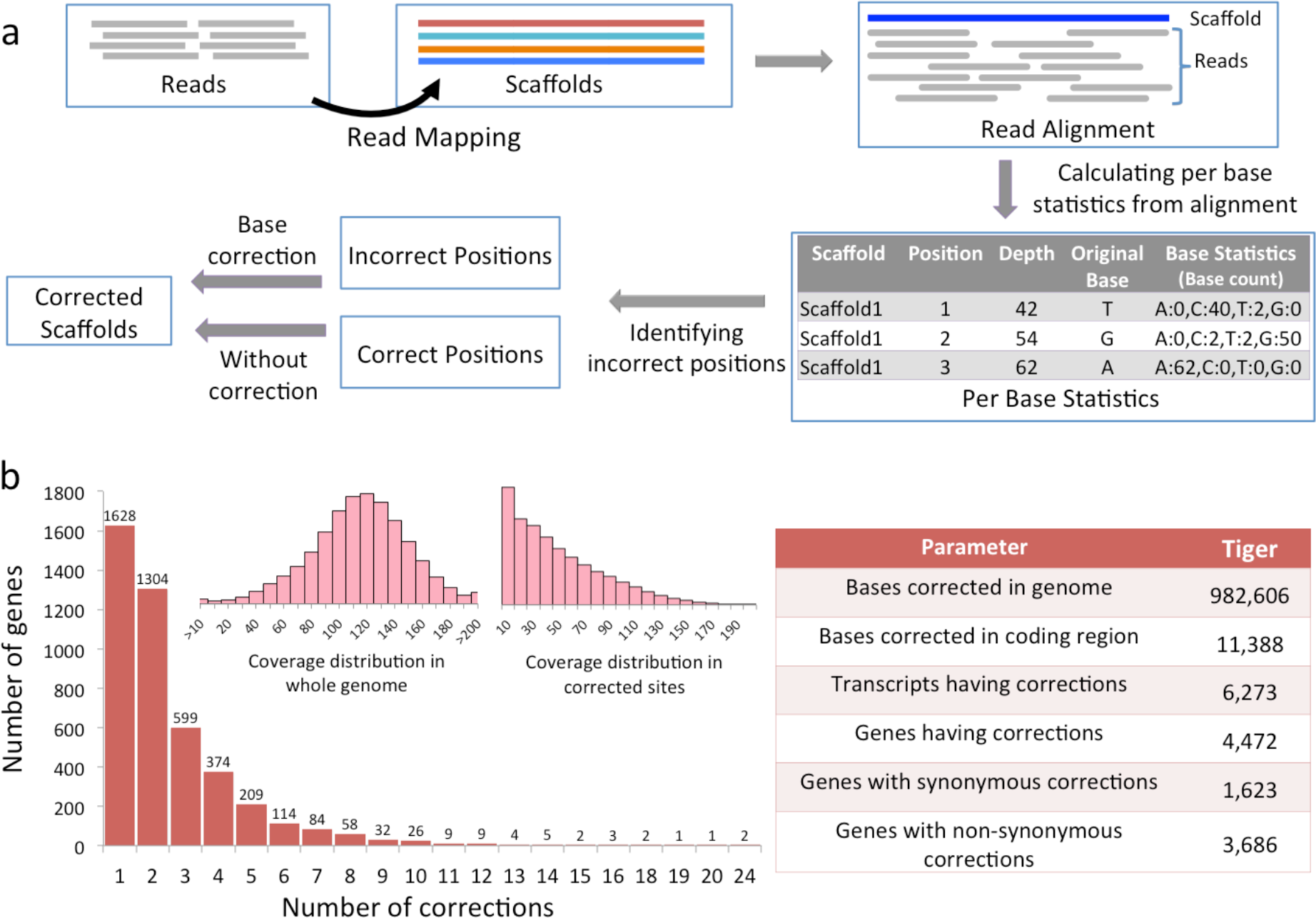
Correction in the tiger genome. **a** Workflow of methods used to identify erroneous sites in the tiger genome assembly and their corrections. **b** The main bar plot represents the number of genes and the number of sites corrected in each gene. The left inset bar plot represents the distribution of coverage of each position in the tiger genome, and the right inset bar plot represents the distribution of coverage of only the corrected sites in the genome assembly. The table in the right represent the correction statistics for the tiger genome assembly.

The corrected genome assembly was used to construct the corrected gene set for tiger as per the gene structure information available at Ensembl release 94. A total of 14,145 codons corresponding to 6,273 transcripts and 4,472 genes were corrected. Of these, 3,686 genes (21%) had non-synonymous, and 1,623 genes (9.3%) had synonymous corrections. Not surprisingly, most of the corrected bases in the coding region of genes in tiger were found identical to the corresponding base present in the cat gene orthologs, which validates the correction methodology.

We also corrected the leopard genome assembly using the same approach. A total of 58,566 bases (0.002% of the genome) were corrected. The corrections in the coding regions mapped to 194 codons in 165 transcripts corresponding to 125 genes, of which 40 genes had synonymous changes and 43 genes had non-synonymous changes. It is apparent that much fewer corrections were made in the leopard genome assembly in comparison to the tiger assembly. This was expected because the leopard genome was sequenced at a very high (~300X) coverage compared to tiger (~100X), and the mapping-based correction was already performed by the authors ^9,39^. Fewer corrections in the leopard genome assembly also indicate that the error correction method used in this study was specific enough to identify and correct only the erroneous positions. The corrected assemblies and gene sets of tiger and leopard were utilized to identify the molecular signatures underlying the adaptive evolution of tiger lineage.

### Adaptive evolution analysis

We performed the adaptive evolution analysis for the lineage leading to tiger using five methods: A) Higher dN/dS using branch model: to identify genes with higher rate of evolution, B) High nucleotide substitution: to identify genes with a high rate of mutation by comparing root-to-tip branch lengths, C) Positive selection using the branch-site model: to identify genes with positive selection in the selected clade, D) Unique substitution with functional impact: to identify genes with unique substitution in the selected clade that have significant impact on the protein function, and E) Positively selected amino acid sites: to identify the positively selected sites in a gene. The analysis was performed using nine mammalian species, including the corrected gene set of leopard and tiger, and the high-quality annotated gene sets of seven species (human, mouse, cow, horse, cat, ferret, dog) retrieved from Ensembl (release 94) ^16^. A total of 10,920 one-to-one orthologs for these nine species were identified using Ensembl BioMart ^20^. The phylogenetic tree for these species was derived using the tree published by Nyakatura et al., 2012 ^40^ by employing the tree subset methodology from the “ape” package of R statistical software ^41^ (**Figure 2a**). The adaptive evolution analysis for the lineage leading to tiger provided several new insights into the felid, Panthera and tiger evolution.

**Figure 2.**
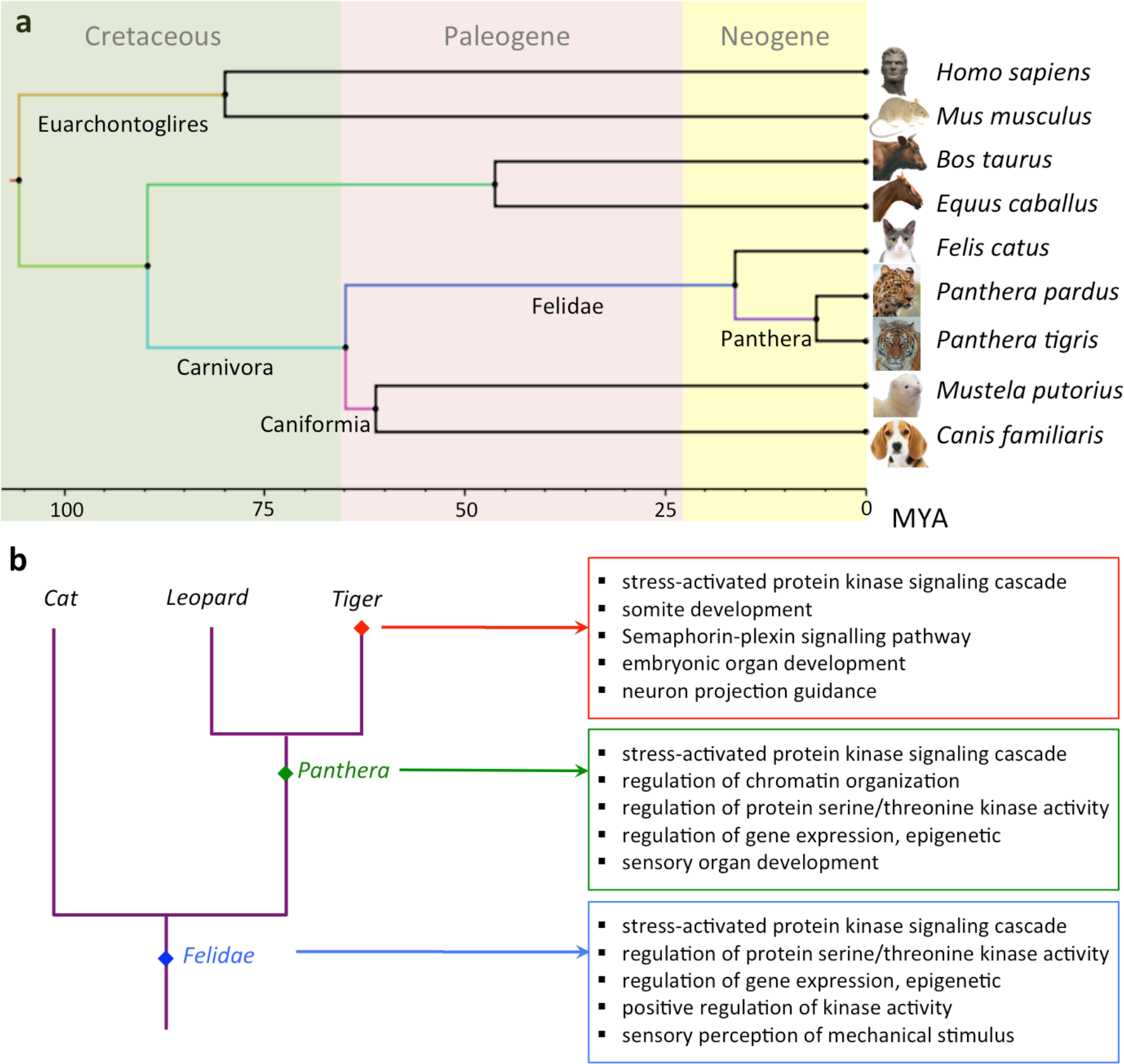
Phylogeny and positive selection in the lineage leading to tiger. **a** The phylogenetic tree of the nine mammalian species used in the study. **b** The top five enriched GO categories of positively selected genes identified in felid, Panthera and tiger lineages.

#### Insights into felid evolution

Recent studies in felids have identified evolutionary signatures that are important for their unique sensory perception and hunting characteristics ^1,5,34,39,42^. However, these studies were performed using the previous gene set (from Cho et. al 2013) of tiger, which contained erroneous base substitutions that can potentially bias the findings. Thus, the usage of corrected tiger gene set in this study is expected to identify the signatures of adaptive evolution in felids. A total of 766 genes showed faster evolution and 906 genes showed positive selection in felid in comparison to the other mammals. These genes showed enrichment for biological functions such as sensory perception, neuronal functioning, cell signalling, development, and stress response (**Figure 2b**). The lists of statistically significant top-20 GO categories from the two analyses are provided in **Supplementary Table S3**. Several genes that previously showed adaptive evolution in felids could not be identified in this study, whereas many additional genes were found to be evolved in felid (**Supplementary Text S2**). However, previous studies on the evolution of felids have also reported positive selection and adaptive evolution in the genes involved in the sensory perception and neuronal functioning ^5^. This indicates that in terms of the broader biological processes, the results from the evolutionary analysis using the corrected genome assemblies corroborate with the previous study on felids ^5^.

We observed several felid-specific amino acid substitutions in the AgRP gene expressed in AGRP neurons, which is involved in regulating the feeding behavior in animals ^43–45^. The injection of AgRP peptides into the brain in rats was found to induce voracious eating behavior even in well-fed mice. AgRP polymorphisms have been associated with diet, leanness, obesity, type-2 diabetes and anorexia nervosa ^45–47^. The felid-specific unique substitutions in the AgRP gene were also found to have significant functional impact predicted using SIFT, and thus, could be associated with the voracious feeding behavior shown by felids ^48,49^.

#### Insights into Panthera evolution

The Panthera genus has shown a recent and rapid diversification, which now comprises of five species of modern big cats possessing several unique characteristics. To understand the genetic-basis of divergence within these species and as well as with respect to the other mammalian species, we performed the comparative evolutionary analysis of Panthera considering tiger and leopard, with seven other mammalian species. The analysis resulted in a total of 1,450 genes showing positive selection in Panthera, which were functionally enriched in sensory perception, regulation of protein serine/threonine kinase activity, gene expression regulation, stress response, and development. A total of 917 genes showed a faster rate of evolution (branch model), and were enriched in cell-cell signalling and early development functions. Further, 797 genes showed amino acid substitutions unique to Panthera with significant functional impact. These genes were enriched in the biological functions related to sperm motility, development, fatty acid metabolism, and DNA repair. The lists of statistically significant top-20 GO categories from the three analyses are provided in **Supplementary Table S4**. A previous study in Panthera had reported unique substitutions with functional impact in fatty acid metabolism and DNA repair categories ^1^, which were also observed in this study.

#### Insights into tiger evolution

A comprehensive analysis of the five types of evolutionary signals was performed using the gene orthologs identified from nine mammalian species to gain insights into the evolution of tiger. A total of 1,474 genes showed positive selection in tiger (branch-site model) and were enriched for functional categories such as early development, fatty acid metabolism and neuronal functioning (**Supplementary Table S5**). A total of 872 genes showed faster evolution in tiger (branch model) and were mainly enriched for functions related to organ development and sensory perception (**Supplementary Table S6**). A total of 1,158 genes showed unique substitutions with functional impact and were enriched for cell signalling, sensory perception, and cytoskeleton functions (**Supplementary Table S7**). A total of 151 genes showed high nucleotide divergence rate identified using root-to-tip branch length values in tiger. These genes were enriched for sensory perception, organ development, and neuronal related functions (**Supplementary Table S8**).

### Insights into the evolution of tiger using genes with multiple signs of adaptation

The genes with multiple signs of adaptation (MSA) were identified as the genes that showed three or more signs of adaptive evolution out of the five methods used for the adaptive evolution analysis (A. Higher dN/dS analysis using the branch model, B. High nucleotide substitution, C. Positive selection using the branch-site model, D. Unique substitution with functional impact, and E. Positively selected amino acid sites). A total of 955 genes showed MSA in tiger in comparison to all the other species including the closely related leopard genome. A total of 83 genes had all the five signs of adaptive evolution, and a maximum of 348 genes showed four signs of adaptive evolution including higher branch dN/dS, positive selection, unique substitution with functional impact, and positively selected sites.

Among the five signatures of adaptive evolution used in this study, the higher branch dN/dS, positive selection, and higher nucleotide divergence can identify the ‘gene-wide’ signals of evolution, suggesting that the complete gene is evolving. On the other hand, unique substitution with functional impact and positively selected sites indicate the evolution of only specific sites in the gene, thus can identify the ‘site-specific’ signals of evolution. Among the MSA, seven genes did not show positive selection (gene-wide signal) in tiger, though they had statistically significant positively selected amino acid sites (site-specific signal), suggesting that the effect of these positively selected sites was masked by the sites evolving under purifying selection or neutrality. Thus, the usage of five different evolutionary analyses helped to identify both site-specific and gene-wide signals of evolution in genes. Using these methods, among the MSA genes, a total of 111 genes showed all the three gene-wide signals of evolution, and a total of 580 genes showed the two site-specific signals of evolution in tiger (**Figure 3a**).

**Figure 3.**
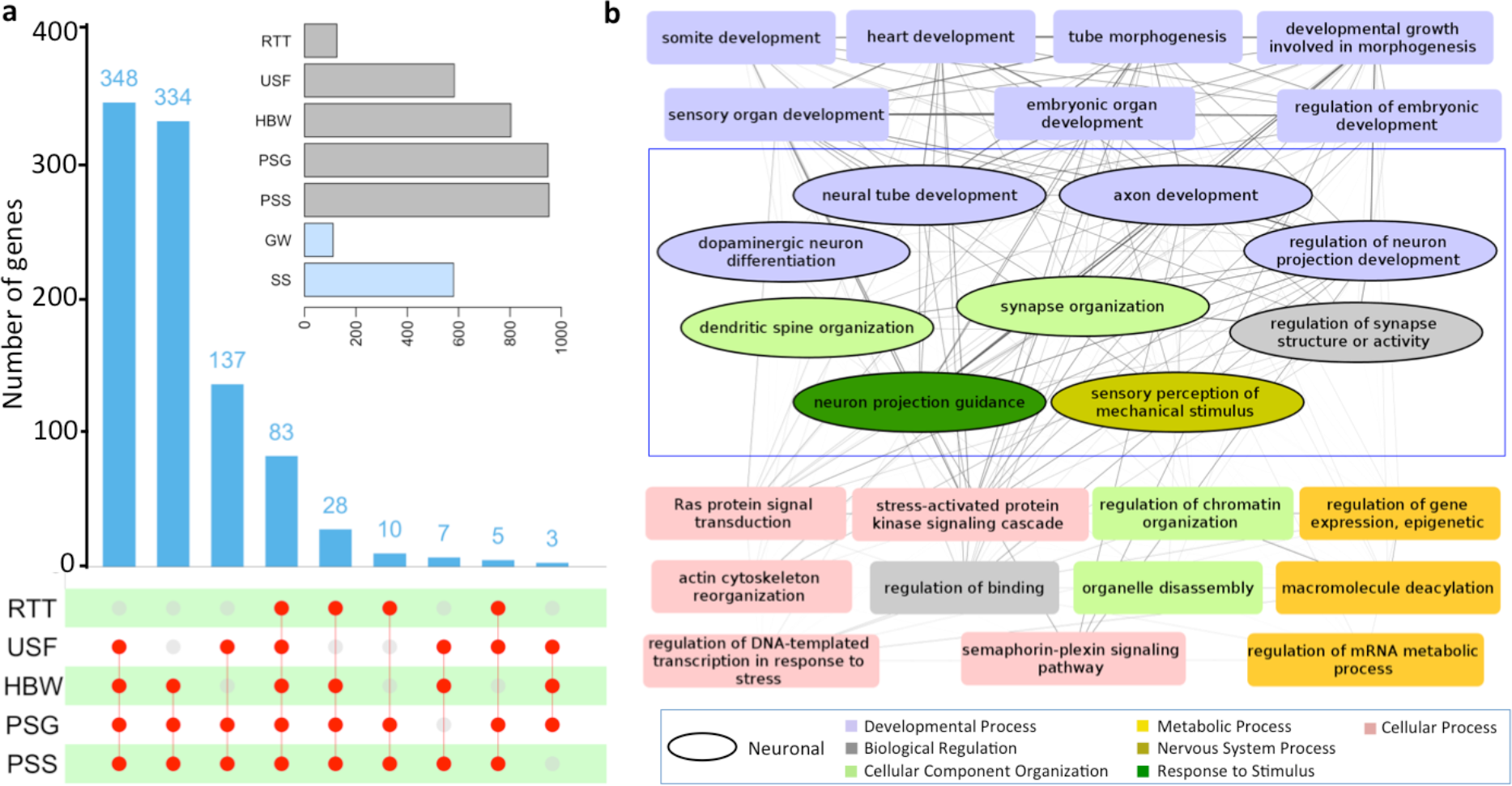
The genes showing multiple signatures of adaptation in tiger. **a** The upSet plot of the number of genes shared by the combination of the five methods used to test for adaptive evolution. The matrix layout was constructed using the upSET package in R ^68^. The connection between the red circles shows the intersection of different methods with the intersection value depicted as a bar plot. RTT: higher root-to-tip branch length, USF: Unique substitution with functional impact, HBW: Higher branch dN/dS (ω), PSG: Positively selected genes, PSS: Positively selected sites, GW: Gene-wide, SS: Site-specific. Gene-wide (GW) represents the genes showing all three signs of adaptation among the MSA (HBW, RTT, PSG), which takes into account the evolution of the complete gene. Site-specific (SS) represents the genes showing the two signs of adaptation among the MSA (USF and PSS), which takes into account the evolution of specific sites in a gene. **b** Network diagram of GO biological processes enriched (p-value < 0.01) in the MSA genes. The nodes represent the GO biological processes, and the edges represent the number of MSA genes shared among the enriched categories.

#### Evolution of developmental and neuronal processes in tiger

The GO enrichment for the MSA genes was performed to identify the underlying biological processes of genes showing adaptive evolution, and the enriched categories (p-value < 0.01) were visualized as a network using Cytoscape v3.2.1 ^50^. The nodes in the network represent the individual GO categories, and the width of edges represents the number of shared genes among the GO categories (**Figure 3b**). It is interesting to note that one-third of the enriched categories were involved in neuronal functioning and development. The genes in these categories perform diverse functions such as regulation, cellular component organization, developmental process, nervous system process, and response to stimulus (**Figure 3b**). It is also apparent from the network that several GO categories, including neuronal-related functions, belonged to a broader GO term “Developmental process”. These GO categories showed dense connections with each other, which indicates that a large number of genes are common among these functional categories. This suggests that these developmental genes with adaptive divergence in tiger have pleiotropic functions, where one gene can regulate multiple developmental processes. Further, the eggNOG classification of the MSA genes revealed ‘signal transduction mechanisms’ as the most enriched category (**Supplementary Table S9**). Taken together, it points towards the differential evolution of neuronal functioning and developmental processes genes in tiger.

#### The highly evolved Notch signalling pathway in tigers

The pathway enrichment analysis performed using the fisher’s exact test and network enrichment method revealed the Notch signalling pathway to be the most significantly enriched pathway (**Supplementary Table S10**). The regression of XD-score, which is a measure of network interconnectivity, and Fisher’s test (with Benjamini-Hochberg adjusted q-value) also revealed that after applying these tests, only the Notch signalling pathway was above the significance threshold (**Figure 4a**). This kind of framework for the pathway enrichment is more accurate than the classical overrepresentation-based method, as it also includes the protein interaction network information ^33^. In the Notch signalling pathway, 11 genes showed adaptive evolution in tiger among which, CTBP1 gene showed all the five signs of adaptation. The 11 genes include the notch receptor (NOTCH3), ligand (DLL3), intracellular and extracellular regulators (DVL3, NUMB, LFNG, ADAM17), transcription factor (RBPJL), and its regulators (CREBBP, NCOR2, CTBP1). From the protein-protein interaction data obtained from STRING database ^51^, it was apparent that these 11 genes can interact with all the genes and regulators of the Notch pathway (**Figure 4b and 4c**). Taken together, it is apparent that every crucial step of the Notch signalling pathway has evolved in tiger in comparison to the other mammalian species, including the close relative leopard. The genes of this pathway are evolutionarily conserved in multi-cellular organisms and regulate the cell-fate determination and tissue homeostasis, thus play an important role in embryonic development ^52,53^. Using the juxtacrine signalling method it regulates the development and functioning of cardiac, neuronal, immune, and endocrine system ^53–55^. The tissue expression data from GNF Atlas ^56^ revealed that these 11 adaptively evolved Notch pathway genes also show high expression in the temporal lobe, whole brain, cerebellum peduncles, and prostate (**Supplementary Table S11**). Thus, the evolution of Notch signalling relates well with the differential neural morphology observed in tiger in comparison to the other mammals ^57,58^.

**Figure 4.**
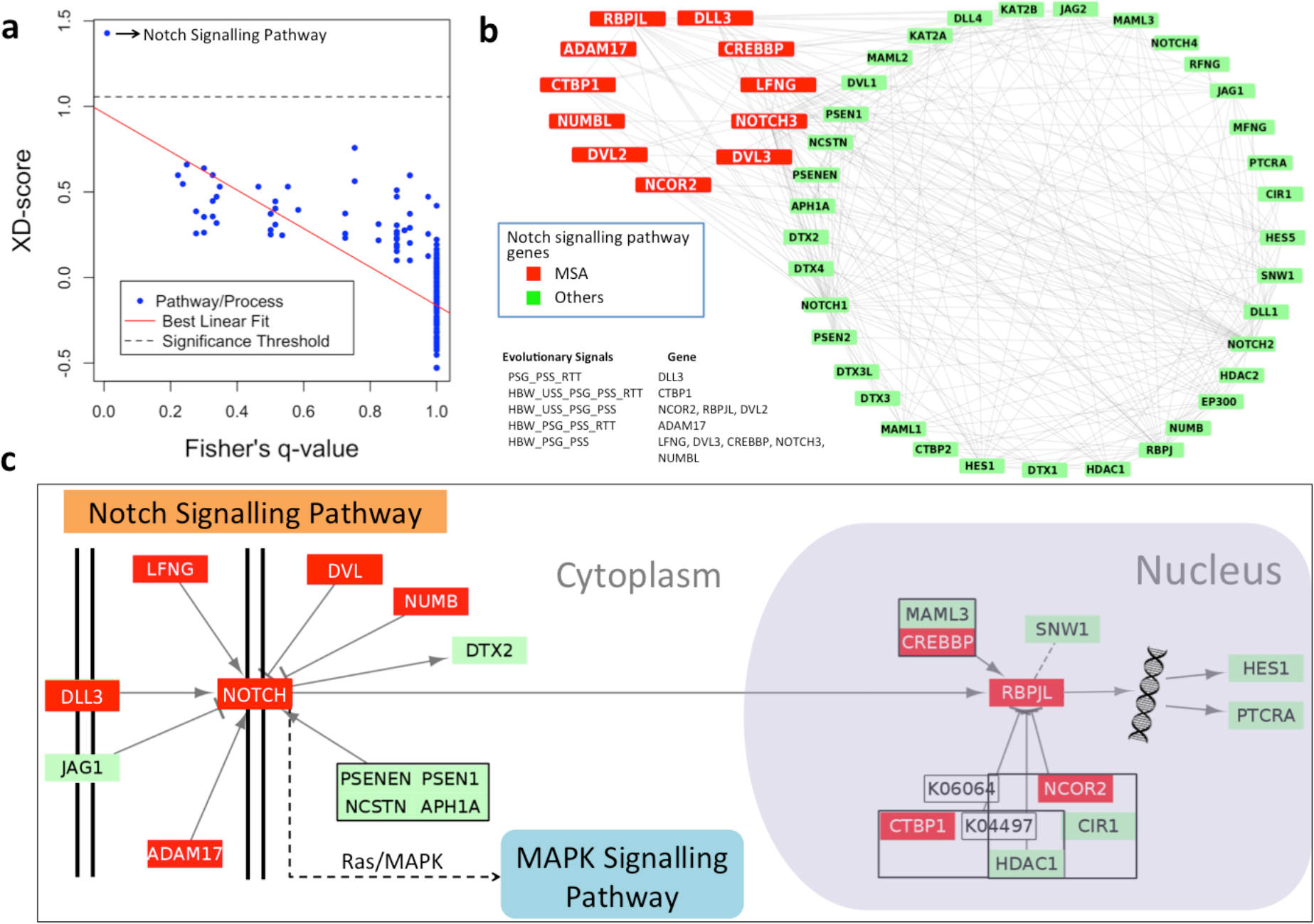
The adaptive divergence of Notch signalling pathway in tiger. **a** The regression of XD-score and Fisher’s q-value of KEGG pathways. **b** Network diagram of genes showing the interaction of MSA genes with the rest of the genes of the Notch signalling pathway. The edges in the network represent the protein-protein interactions among the genes obtained from STRING database. **c** Schematic representation of the Notch signalling pathway from KEGG with the genes showing multiple signatures of adaptation highlighted in Red. The pathway diagram was constructed using KEGGscape plug-in ^69^ in Cytoscape.

## DISCUSSION

Genome sequencing followed by genome-wide comparative analysis has become a powerful tool to study the patterns of evolution in different lineages. The genome sequencing of tiger has provided novel insights into their unique adaptations and divergence from other species, and among its subspecies. The genome sequencing of tiger is significant since it is a part of charismatic megafauna that has captivated human interest, is the largest felid, and is among one of the most endangered species with less than 4,000 individuals remaining in the wild ^59^.

In this study, while using the publicly available genome sequence assembly of tiger, we found that the assembly consisted of several errors, which also led to several incorrect interpretations in recent other studies. For example, Figueiro et al., 2017 identified that the ESRP1 gene, important for craniofacial robustness, has a positively selected I298Y substitution in jaguar ^34^. This substitution was found to be positively selected in jaguar due to the presence of “I” in the respective ortholog in tiger, which was a result of single nucleotide error in the tiger genome assembly (**Supplementary Figure 3 and 4**). From the above example and the other cases described in the **Supplementary Text**, it is apparent that evolutionary studies are very sensitive to nucleotide errors present in the genome assemblies and gene sets, where even single nucleotide errors can produce drastically misleading results in the analyses.

Thus, to understand the adaptive evolution of the lineage leading to tiger, the publicly available tiger genome assembly was corrected for such single nucleotide errors using the ‘SeqBug’ pipeline developed in this study and validated using the resequencing data of a new male Bengal tiger genome and transcriptome. The identification of errors in 4,472 genes and 982,606 bases (0.04% genome) in the tiger genome put forth the need of the correction. Further, the incorrect positions were mostly present in regions of low coverage (<30) and were much fewer in the leopard genome that was sequenced at three times higher coverage than tiger ^9,39^. These observations underscore the need for a higher genome coverage along with mapping-based correction to produce a more accurate assembly. The genome sequence of another tiger individual sequenced in this study and the construction of corrected genome sequence and gene set of tiger are likely to be beneficial for further comparative studies.

After correction, most of the bases corrected in the coding genome of tiger were identical to the corresponding bases in the cat genome, thus, validating our correction methodology. This further indicates that the divergence time of tiger calculated using the genetic differences in the previous studies could suffer from an over-estimation because of these erroneous substitutions ^8^. Considering errors of 0.9 million bases and mutation rate of 1.1e-09 per base per year for tiger ^1^, the estimated divergence time of tiger can be affected by 0.37 million years.

The usage of corrected tiger and leopard coding genome, a large number of orthologs, and five different evolutionary analysis in this study, makes the evolutionary assessments more reliable, and was also successful in revealing the signatures of adaptive divergence in felids, Panthera and tiger lineages. Previous reports identified evolutionary adaptations in genes related to muscle strength, hypercarnivorous diet, sensory perception, and craniofacial and limb development in the Panthera/Felidae lineage ^1,5,34,39^. Similar categories were also found to be adaptively evolved in the Panthera/Felidae lineage in this study. However, large differences in the gene sets were observed, which further highlights the impact of incorrect genomes on the evolutionary analysis.

One of the unique finding was the enrichment of neuronal functioning and developmental processes in genes showing multiple signs of adaptive evolution in tiger. Notably, the Notch signalling pathway emerged as the most diverged pathway in tiger, which was not found as adaptively evolved in the previous studies. The observation is significant since the Notch pathway plays key roles in diverse developmental processes including neurogenesis, neural differentiation, and cell fate determination ^53,60^. Also, the observed divergence at almost every step of Notch signalling pathway, which is a highly conserved pathway throughout the animal kingdom, further indicates the adaptive evolution in neuronal functioning and development genes in tiger.

The evolution of the neuronal related genes in the tiger lineage is informative but not very surprising given their unique physiological and behavioral characteristics ^2,3,49,61–63^. Several studies show that the feeding, drinking, aggression, predation and sexual behavior, and the energy homeostasis of an organism are primarily governed by neuronal circuitry ^43,44,64–66^. The felids, particularly the big cats being the large hypercarnivores, show a very distinct aggressive and predatory behavior. They require strong neuro-muscular coordination, sensory perception and timed actions for successful hunting ^49,67^. Thus, it is tempting to speculate the role of evolution in the neural development and processes for attaining unique phenotypes, behavior, and dietary patterns. This notion also gets support from the previous studies that showed differences in neuronal morphology in tiger in comparison to other mammals, including its closest relative leopard ^57,58^. The dendrites of typical pyramidal neurons in tiger are very complex, and the dendritic measures of these neurons are disproportionally large relative to body/brain size ^58^. To summarize, the identification of adaptive evolution in the neuronal functioning genes in tiger indicates the plausible role of evolution in neural processes in achieving exceptional sensory perception, neuro-muscular coordination, faster reflex actions, predatory capabilities and hypercarnivorous behavior in tiger.

## DATA ACCESSIBILITY

The sequence data of the Bengal tiger genome and transcriptome will be made publicly available on acceptance of the manuscript. The corrected genome assemblies, gene sets, and correction pipeline ‘SeqBug’ developed in this study are available from the corresponding author on reasonable request.

**LIST OF ABBREVIATIONS**
USF: Unique substitutions with functional impact
PSG: Postively selected genes
PSS: Postively selected sites
HBW: Higher branch dN/dS
BEX3: Brain expressed gene 3
AgRP: Agouti related neuropeptide
ESRP1: Epithelial splicing regulatory protein1

## ACKNOWLEDGMENT

We thank Dr. Atul Gupta, Wildlife Veterinary Officer, Van Vihar National Park, Bhopal, and Director, Van Vihar National Park, Bhopal, India for providing the blood samples of tiger. We also acknowledge the help of Drs. Tista Joseph and Niraj Dahe, Wildlife Veterinary Officers (Wildlife SOS India) at Van Vihar National Park for carrying out the sample collection procedure. We thank the HPC facility and NGS facility at IISER Bhopal. SKJ and RS thank the Department of Science and Technology for the DST-INSPIRE fellowship. We also thank the intramural research funds provided by IISER Bhopal.

## AUTHOR’S CONTRIBUTION

VKS conceived and coordinated the project. RS prepared the DNA samples and performed genome sequencing. PM and SKJ generated the corrected genome assemblies and gene sets. PM performed the branch dN/dS, positive selection, unique substitution and SIFT analyses. SKJ performed the higher nucleotide divergence analysis. PM, SKJ, NV, and VKS analyzed the data and wrote the manuscript. PM created figures. All the authors have read and approved the final manuscript.

