## Supplementary Material for "Comparative analysis of corrected tiger genome provides clues to their neuronal evolution"

*Corresponding author email:

Email address of authors:

### Supplementary Figures

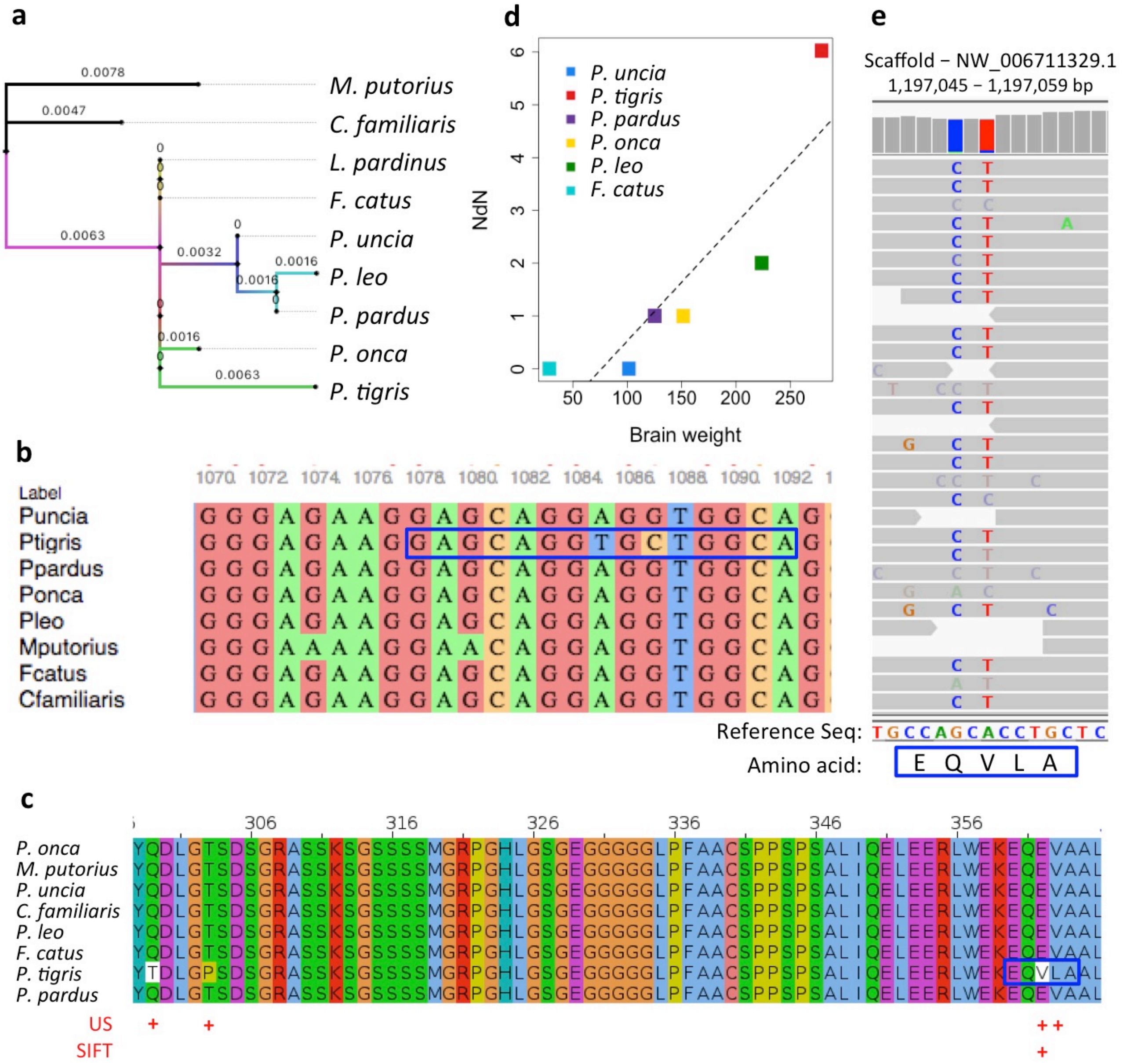


**Supplementary Figure S1. Multiple signs of adaptation in LZTS3 gene using the incorrect tiger geneset. a** higher branch length, **b** nucleotide alignment, **c** protein alignment showing unique substitutions (US) and substitution with functional impact (SIFT), **d** phylogenetically corrected correlation of LZTS3 gene with brain weight, **e** visualization of the site identified as having unique substitutions and substitution with functional impact in the gene in IGV. Two unique substitutions VL, with V as SIFT substitution in the blue box have been shown in the IGV along with its neighboring sites. The gene exists on the (-) strand of the genome and its (+) strand is shown in **e**.

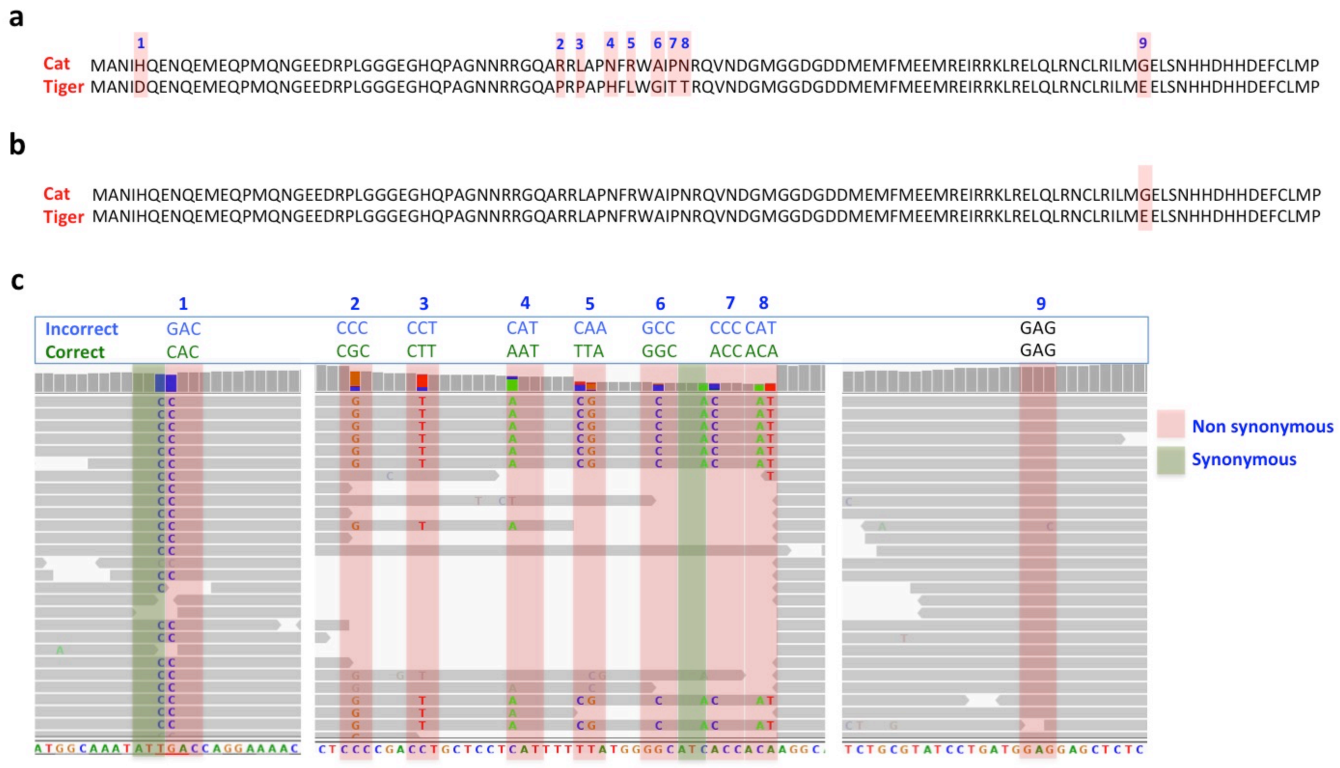


**Supplementary Figure S2: The BEX3 gene in tiger before and after correction. a** The unique amino acid substitutions in BEX3 gene in tiger in the incorrect version. **b** The unique amino acid substitutions in BEX3 gene in tiger in the corrected version. **c** Visualization of the nine substitutions (from incorrect version) in IGV. The sequence alignment has been taken from Ensembl release 94.

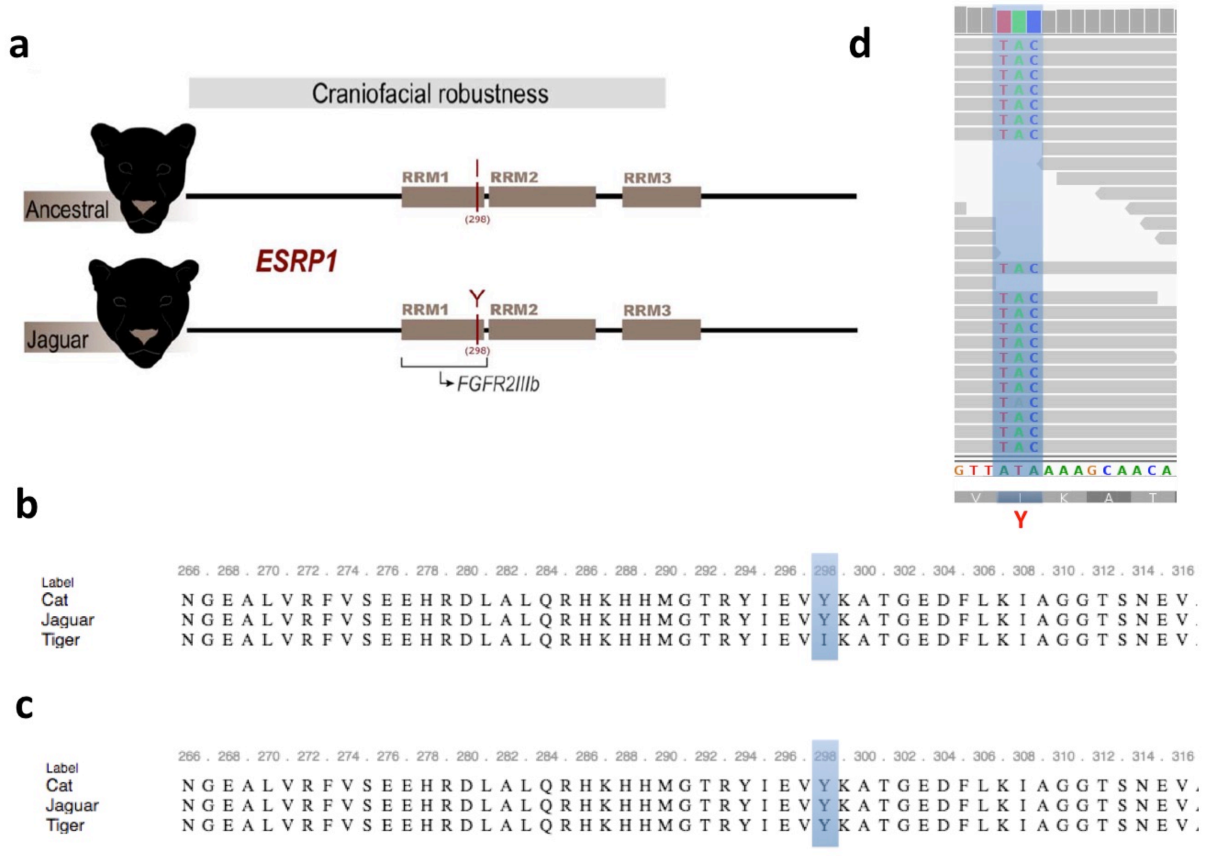


**Supplementary Figure S3. The divergence in ESRP1 gene involved in craniofacial robustness (Correction to Figueiro et al., 2017). a** I298Y mutation in jaguar as identified by Figueiro et al., 2017 **b** The protein alignment for ESRP1 gene showing 298 position in felids using uncorrected tiger assembly. **c** The protein alignment for ESRP1 gene showing 298 position in felids using the corrected tiger assembly. **d** Reads alignment of tiger genome showing 298 position as correctly**.**

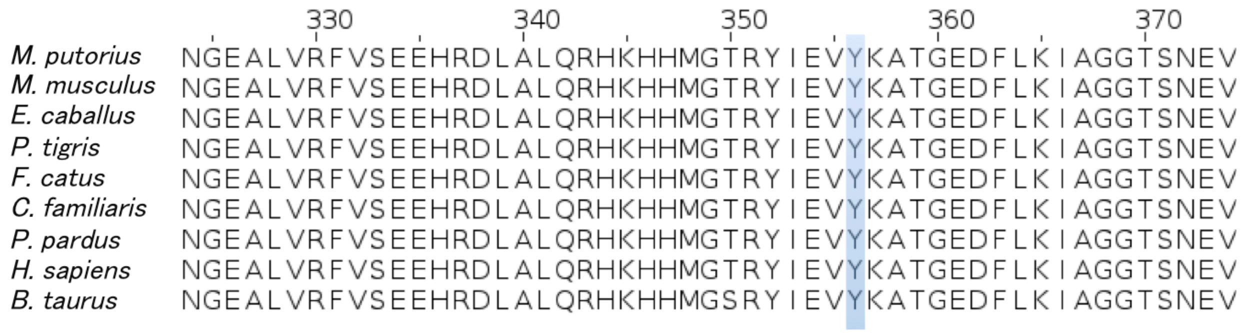


**Supplementary Figure S4. The protein alignment for ESRP1 gene showing 298 position in nine mammalian species (Correction to Figueiro et al., 2017).** An insertion of 58 amino acids at the beginning in *M. putorious* led to a shift in the alignment position to 356.

```
Mmusculus       MLTAMLLSCVLLLALPPTLGVQMGVAPLKGIRRPDQALFPEFPGLSLNG-LKKTTADRAE   59
Hsapiens        MLTAAVLSCALLLALPATRGAQMGLAPMEGIRRPDQALLPELPGLGLRAPLKKTTAEQAE   60
Ecaballus       MLTAVLLSCALLLTLPAMQGAQMGLVPLEGIRT-DQALIPELPGLGIRPLLKKTTAEQAE   59
Btaurus         MLTAVLLSCALLLAMPPMQGAQMGPAPLEGIGRPEEALFLELQGLSLQPSLKRITEEQAE   60
Ptigris         MLTAVLLSWALLLTLPPMQEAQRGLAPLEGIRRPDQALFPELPGLGRQPPLKRTSAEQSI   60
Ppardus         MLTAVLLSWALLLTLPPMQEAQRGLAPLEGIRRPDQALFPELPGLGRQPPLKRTSAEQSI   60
Fcatus          MLTAVLLSWALLLTLPPMQEAQRGLAPLEGIRRPDQALFPELPGLGQQPPLKRTSAEQSI   60
Mputorius       MLPAVLLSCALLLALPPMQGAGIGLAPLEGIRSPDQALFPELPGLGLQSSLKRTAAEQSG   60
Cfamiliaris     MLPAVLLSCALLLALPPMQGAQIGLVPLEGIRRPDQALFPELPGLGLQSSLKRATAEQSE   60
            USF         +          +

Mmusculus       EVLLQKAE--ALAEVLDPQNRESRSPRRCVRLHESCLGQQVPCCDPCATCYCRFFNAFCY   117
Hsapiens        EDLLQEAQ--ALAEVLDLQDREPRSSRRCVRLHESCLGQQVPCCDPCATCYCRFFNAFCY   118
Ecaballus       EALKQEAK--ALAEVLDLEGREQRSPRRCVRLHESCLGHQVPCCDPCATCYCRFFNAFCY   117
Btaurus         ESLLQEAEAKALAEVLDPEGRKPRSPRRCVRLHESCLGHQVPCCDPCATCYCRFFNAFCY   120
Ptigris         EALLQEAE--ALTEVLDPEGREPRSPRRCVRLQESCLGHQVSCCDPCATCYCRFFNAFCY   118
Ppardus         EALLQEAE--ALTEVLDPEGREPRSPRRCVRLQESCLGHQVSCCDPCATCYCRFFNAFCY   118
Fcatus          EALLQEAE--ALTEVLDPEGREPRSPRRCVRLQESCLGHQVSCCDPCATCYCRFFNAFCY   118
Mputorius       EALLQEAE--ALSEVLDPEGRESRSPRRCVRLHESCLGHQVPCCDPCATCYCRFFNAFCY   118
Cfamiliaris     EALLQEAE--ALAEVLDPEGREPRSPRRCVRLHESCLGHQVPCCDPCATCYCRFFNAFCY   118
            USF                                 +

Mmusculus       CRKLGTATNLCSRTX        132
Hsapiens        CRKLGTAMNPCSRTX        133
Ecaballus       CRKLGTAMNPCSRTX        132
Btaurus         CRKLGTTTNPCSRTX        135
Ptigris         CRKLGTATIPCSRTX        133
Ppardus         CRKLGTATIPCSRTX        133
Fcatus          CRKLGTATIPCSRTX        133
Mputorius       CRKLGTAMNPCSRTX        133
Cfamiliaris     CRKLGTATNPCSRT-        132
            USF         +
```

**Supplementary Figure S5. Alignment of AgRP gene orthologs from nine mammalian species showing felid specific amino acid substitutions.** The sites with unique substitution with functional impact (USF) in felids have been indicated with "+" sign.

**Supplementary Tables**

**Supplementary Table S1: Summary of genome sequencing of Bengal tiger.**

| Library Name | Library Preparation method | Insert Size | Sequencer | Read length | Raw reads |
|---|---|---|---|---|---|
| TG-1 | Nextera XT | 650 bp | NextSeq 500 | 150*2 | 21,003,653 |
| TG-2 | Nextera XT | 650 bp | NextSeq 500 | 150*2 | 27,243,831 |
| TG-3 | Truseq PCR free | 550 bp | NextSeq 500 | 150*2 | 52,406,157 |
| CTG | Truseq PCR free | 350bp | HiSeq | 250*2 | 75,027,209 |

**Supplementary Table S2: Summary of transcriptome sequencing of Bengal tiger.**

| | Library preparation method | Sequencer | Raw reads |
|---|---|---|---|
| Library 1 | TruSeq RNA Sample prep Kit | HiSeq 2500 | 15,823,343 |
| Library 2 | TruSeq RNA Sample prep Kit | HiSeq 2500 | 16,429,561 |

**Supplementary Table S3: Adaptive evolution in felids. The top 20 GOs identified as enriched in positive selection and higher branch dN/dS (HBW) analyses in felids after correction.**

| Positive selection | | | HBW | | |
|---|---|---|---|---|---|
| Description | O | p-value | Description | O | p-value |
| stress-activated protein kinase signaling cascade | 30 | 0.0001 | sensory perception of mechanical stimulus | 21 | 0.0001 |
| regulation of protein serine/threonine kinase activity | 47 | 0.0001 | synapse organization | 23 | 0.0005 |
| regulation of gene expression, epigenetic | 25 | 0.0003 | establishment or maintenance of cell polarity | 18 | 0.0007 |
| positive regulation of kinase activity | 48 | 0.0004 | organelle localization | 39 | 0.0012 |
| sensory perception of mechanical stimulus | 21 | 0.0007 | muscle cell differentiation | 31 | 0.0019 |
| organ or tissue specific immune response | 5 | 0.0007 | vesicle-mediated transport in synapse | 14 | 0.0019 |
| neuron projection guidance | 24 | 0.0016 | multi-organism metabolic process | 15 | 0.0020 |
| tumor necrosis factor superfamily cytokine production | 13 | 0.0018 | morphogenesis of an epithelial sheet | 8 | 0.0025 |
| sensory organ development | 46 | 0.0025 | sensory organ development | 41 | 0.0026 |
| gene silencing | 22 | 0.0025 | regulation of binding | 26 | 0.0026 |
| axon development | 40 | 0.0030 | mesenchyme development | 22 | 0.0030 |
| cell-cell signaling by wnt | 40 | 0.0040 | stress-activated protein kinase signaling cascade | 23 | 0.0032 |
| maintenance of location | 26 | 0.0043 | neurotransmitter transport | 19 | 0.0034 |

| | | | | | |
|---|---|---|---|---|---|
| regulation of intracellular transport | 40 | 0.0062 | respiratory system development | 20 | 0.0037 |
| embryonic organ development | 36 | 0.0082 | response to inorganic substance | 36 | 0.0053 |
| regulation of binding | 27 | 0.0086 | regulation of neurotransmitter levels | 18 | 0.0070 |
| tube morphogenesis | 31 | 0.0096 | developmental growth involved in morphogenesis | 20 | 0.0074 |
| beta-catenin-TCF complex assembly | 6 | 0.0098 | urogenital system development | 27 | 0.0075 |
| inclusion body assembly | 4 | 0.0099 | connective tissue development | 21 | 0.0077 |
| syncytium formation | 8 | 0.0100 | trabecula morphogenesis | 8 | 0.0084 |

**Supplementary Table S4: Adaptive evolution in Panthera. The top 20 GOs identified as enriched in positive selection, unique substitution with functional impact (USF) and higher branch dN/dS (HBW) analyses in Panthera after correction.**

| Positive selection | | | USF | | | HBW | | |
|---|---|---|---|---|---|---|---|---|
| **Description** | **O** | **p-value** | **Description** | **O** | **p-value** | **Description** | **O** | **p-value** |
| stress-activated protein kinase signaling cascade | 43 | 0.0000 | sperm motility | 12 | 0.0000 | tube morphogenesis | 39 | 0.0001 |
| regulation of chromatin organization | 25 | 0.0020 | cilium organization | 25 | 0.0017 | developmental growth involved in morphogenesis | 27 | 0.0004 |
| regulation of protein serine/threonine kinase activity | 62 | 0.0022 | ventricular system development | 5 | 0.0092 | cell-cell signaling by wnt | 45 | 0.0005 |
| regulation of gene expression, epigenetic | 32 | 0.0022 | regulation of response to wounding | 14 | 0.0114 | regulation of chromatin organization | 20 | 0.0005 |
| sensory organ development | 68 | 0.0022 | protein complex localization | 10 | 0.0132 | endocrine system development | 16 | 0.0006 |
| sensory perception of mechanical stimulus | 26 | 0.0068 | necrotic cell death | 6 | 0.0150 | sensory perception of mechanical stimulus | 22 | 0.0008 |
| pattern specification process | 47 | 0.0114 | neurotrophin signaling pathway | 5 | 0.0211 | Ras protein signal transduction | 35 | 0.0010 |
| dendritic spine organization | 10 | 0.0116 | lipid catabolic process | 22 | 0.0242 | stress-activated protein kinase signaling cascade | 28 | 0.0011 |
| embryonic organ development | 52 | 0.0132 | microtubule-based movement | 17 | 0.0277 | regulation of binding | 31 | 0.0012 |
| cell-cell signaling by wnt | 56 | 0.0134 | DNA repair | 34 | 0.0284 | synapse organization | 25 | 0.0013 |
| mesenchyme development | 32 | 0.0140 | cell-substrate adhesion | 24 | 0.0285 | connective tissue development | 27 | 0.0014 |
| protein maturation | 31 | 0.0169 | peptide cross-linking | 4 | 0.0292 | somite development | 13 | 0.0014 |
| neuron projection guidance | 30 | 0.0171 | sodium ion transport | 18 | 0.0294 | peptidyl-serine modification | 30 | 0.0017 |
| semaphorin-plexin signaling pathway | 8 | 0.0175 | neutral lipid metabolic process | 11 | 0.0310 | vesicle-mediated transport in synapse | 16 | 0.0018 |
| tube morphogenesis | 44 | 0.0187 | fatty acid metabolic | 24 | 0.0327 | regulation of protein | 44 | 0.0021 |

| | | | | | | | | |
|---|---|---|---|---|---|---|---|---|
| | | | process | | | serine/threonine kinase activity | | |
| somite development | 14 | 0.0194 | digestion | 11 | 0.0335 | gene silencing | 23 | 0.0021 |
| macromolecule deacylation | 14 | 0.0194 | skin development | 16 | 0.0350 | regulation of morphogenesis of an epithelium | 22 | 0.0023 |
| developmental growth involved in morphogenesis | 30 | 0.0203 | lipid homeostasis | 11 | 0.0361 | neural tube development | 19 | 0.0025 |
| skeletal system development | 57 | 0.0209 | response to nerve growth factor | 5 | 0.0400 | muscle cell differentiation | 35 | 0.0028 |
| insulin-like growth factor receptor signaling pathway | 8 | 0.0220 | interleukin-8 production | 6 | 0.0404 | respiratory system development | 23 | 0.0028 |

**Supplementary Table S5: The top 20 GO identified as enriched in genes of positively selected genes in tiger after correction.**

| Geneset | Description | C | O | p-value |
|---|---|---|---|---|
| GO:0031098 | stress-activated protein kinase signaling cascade | 172 | 42 | 0.0001 |
| GO:0061053 | somite development | 58 | 16 | 0.0046 |
| GO:0071526 | semaphorin-plexin signaling pathway | 26 | 9 | 0.0064 |
| GO:0048568 | embryonic organ development | 286 | 55 | 0.0068 |
| GO:0097485 | neuron projection guidance | 151 | 32 | 0.0087 |
| GO:0071774 | response to fibroblast growth factor | 95 | 22 | 0.0098 |
| GO:0045995 | regulation of embryonic development | 84 | 20 | 0.0098 |
| GO:0050954 | sensory perception of mechanical stimulus | 118 | 26 | 0.0103 |
| GO:0198738 | cell-cell signaling by wnt | 312 | 58 | 0.0113 |
| GO:0040029 | regulation of gene expression, epigenetic | 143 | 30 | 0.0125 |
| GO:0007389 | pattern specification process | 252 | 48 | 0.0130 |
| GO:0097061 | dendritic spine organization | 34 | 10 | 0.0144 |
| GO:0030323 | respiratory tube development | 127 | 27 | 0.0145 |
| GO:0022604 | regulation of cell morphogenesis | 305 | 56 | 0.0160 |
| GO:0061564 | axon development | 307 | 56 | 0.0182 |
| GO:0006631 | fatty acid metabolic process | 226 | 43 | 0.0183 |
| GO:0030900 | forebrain development | 235 | 44 | 0.0223 |
| GO:0001101 | response to acid chemical | 223 | 42 | 0.0230 |
| GO:0007423 | sensory organ development | 362 | 64 | 0.0230 |
| GO:0071900 | regulation of protein serine/threonine kinase activity | 324 | 58 | 0.0234 |

**Supplementary Table S6: The top 20 GO identified as enriched in genes showing accelerated rate of evolution in tiger (as determined by higher branch dN/dS) after correction.**

| Geneset | Description | C | O | p-value |
|---|---|---|---|---|
| GO:0061053 | somite development | 54 | 15 | 0.0001 |
| GO:0007507 | heart development | 332 | 51 | 0.0001 |
| GO:0007423 | sensory organ development | 339 | 51 | 0.0002 |
| GO:0050954 | sensory perception of mechanical stimulus | 108 | 22 | 0.0002 |

| | | | | |
|---|---|---|---|---|
| GO:0031098 | stress-activated protein kinase signaling cascade | 158 | 28 | 0.0004 |
| GO:0060560 | developmental growth involved in morphogenesis | 145 | 26 | 0.0006 |
| GO:0097485 | neuron projection guidance | 142 | 25 | 0.0010 |
| GO:1902275 | regulation of chromatin organization | 97 | 19 | 0.0010 |
| GO:1903311 | regulation of mRNA metabolic process | 76 | 16 | 0.0011 |
| GO:0040029 | regulation of gene expression, epigenetic | 137 | 24 | 0.0013 |
| GO:0198738 | cell-cell signaling by wnt | 289 | 42 | 0.0015 |
| GO:0030900 | forebrain development | 221 | 34 | 0.0016 |
| GO:0035239 | tube morphogenesis | 230 | 35 | 0.0017 |
| GO:0016458 | gene silencing | 127 | 22 | 0.0023 |
| GO:0061564 | axon development | 287 | 41 | 0.0024 |
| GO:0051098 | regulation of binding | 186 | 29 | 0.0028 |
| GO:0071900 | regulation of protein serine/threonine kinase activity | 299 | 42 | 0.0029 |
| GO:0007265 | Ras protein signal transduction | 221 | 33 | 0.0030 |
| GO:0050803 | regulation of synapse structure or activity | 69 | 14 | 0.0033 |
| GO:1903008 | organelle disassembly | 69 | 14 | 0.0033 |

**Supplementary Table S7: GO enrichment of genes showing unique substitution with functional impact in tiger after correction.**

| Geneset | Description | C | O | p-value |
|---|---|---|---|---|
| GO:0034067 | protein localization to Golgi apparatus | 20 | 8 | 0.0007 |
| GO:0050954 | sensory perception of mechanical stimulus | 119 | 22 | 0.0072 |
| GO:0071774 | response to fibroblast growth factor | 95 | 18 | 0.0112 |
| GO:0008202 | steroid metabolic process | 166 | 27 | 0.0174 |
| GO:0016358 | dendrite development | 137 | 23 | 0.0190 |
| GO:0051493 | regulation of cytoskeleton organization | 279 | 41 | 0.0216 |
| GO:0009410 | response to xenobiotic stimulus | 40 | 9 | 0.0230 |
| GO:0097061 | dendritic spine organization | 34 | 8 | 0.0243 |
| GO:0022604 | regulation of cell morphogenesis | 306 | 44 | 0.0250 |
| GO:0032886 | regulation of microtubule-based process | 96 | 17 | 0.0251 |
| GO:0044782 | cilium organization | 187 | 29 | 0.0256 |
| GO:0070849 | response to epidermal growth factor | 23 | 6 | 0.0304 |
| GO:0043062 | extracellular structure organization | 230 | 34 | 0.0317 |
| GO:0023019 | signal transduction involved in regulation of gene expression | 12 | 4 | 0.0320 |
| GO:0000226 | microtubule cytoskeleton organization | 295 | 42 | 0.0324 |
| GO:0050906 | detection of stimulus involved in sensory perception | 70 | 13 | 0.0330 |
| GO:0045730 | respiratory burst | 18 | 5 | 0.0365 |
| GO:0006631 | fatty acid metabolic process | 226 | 33 | 0.0395 |
| GO:0050951 | sensory perception of temperature stimulus | 13 | 4 | 0.0425 |
| GO:0051604 | protein maturation | 157 | 24 | 0.0458 |
| GO:0051235 | maintenance of location | 181 | 27 | 0.0466 |
| GO:0050953 | sensory perception of light stimulus | 142 | 22 | 0.0478 |

**Supplementary Table S8: The top 20 GO identified as enriched in genes showing higher nucleotide divergence in tiger (as determined by root-to-tip branch length method) after correction.**

| Geneset | Description | C | O | p-value |
|---|---|---|---|---|
| GO:0048483 | autonomic nervous system development | 24 | 4 | 0.0004 |
| GO:0007423 | sensory organ development | 364 | 13 | 0.0025 |
| GO:0060485 | mesenchyme development | 164 | 8 | 0.0028 |
| GO:0061383 | trabecula morphogenesis | 41 | 4 | 0.0029 |
| GO:0021953 | central nervous system neuron differentiation | 108 | 6 | 0.0050 |
| GO:0051962 | positive regulation of nervous system development | 307 | 11 | 0.0053 |
| GO:0061564 | axon development | 309 | 11 | 0.0056 |
| GO:0071542 | dopaminergic neuron differentiation | 25 | 3 | 0.0056 |
| GO:1904888 | cranial skeletal system development | 49 | 4 | 0.0056 |
| GO:0097485 | neuron projection guidance | 152 | 7 | 0.0069 |
| GO:0001764 | neuron migration | 83 | 5 | 0.0073 |
| GO:0050954 | sensory perception of mechanical stimulus | 119 | 6 | 0.0079 |
| GO:0045787 | positive regulation of cell cycle | 201 | 8 | 0.0093 |
| GO:0021510 | spinal cord development | 59 | 4 | 0.0108 |
| GO:0061053 | somite development | 59 | 4 | 0.0108 |
| GO:0007389 | pattern specification process | 253 | 9 | 0.0119 |
| GO:0008637 | apoptotic mitochondrial changes | 64 | 4 | 0.0142 |
| GO:0051961 | negative regulation of nervous system development | 176 | 7 | 0.0147 |
| GO:0035265 | organ growth | 101 | 5 | 0.0162 |
| GO:0090287 | regulation of cellular response to growth factor stimulus | 143 | 6 | 0.0184 |

**Supplementary Table S9: eggNOG classification of MSA genes in tiger after correction.**

| Category Code | eggNOG Category Description | Broad category | Number of genes |
|---|---|---|---|
| [T] | Signal transduction mechanisms | Cellular processes and signaling | 219 |
| [K] | Transcription | Information storage and processing | 138 |
| [S] | Function unknown | Poorly characterized | 99 |
| [O] | Post-translational modification, protein turnover, and chaperones | Cellular processes and signaling | 76 |
| [U] | Intracellular trafficking, secretion, and vesicular transport | Cellular processes and signaling | 64 |
| [Z] | Cytoskeleton | Cellular processes and signaling | 64 |
| [W] | Extracellular structures | Cellular processes and signaling | 52 |
| [P] | Inorganic ion transport and metabolism | Metabolism | 51 |
| [B] | Chromatin structure and dynamics | Information storage and processing | 37 |
| [I] | Lipid transport and metabolism | Metabolism | 28 |
| [A] | RNA processing and modification | Information storage and processing | 27 |
| [G] | Carbohydrate transport and metabolism | Metabolism | 26 |
| [C] | Energy production and conversion | Metabolism | 16 |
| [D] | Cell cycle control, cell division, chromosome partitioning | Cellular processes and signaling | 12 |
| [E] | Amino acid transport and metabolism | Metabolism | 12 |

| | | | |
|---|---|---|---|
| [L] | Replication, recombination and repair | Information storage and processing | 11 |
| [F] | Nucleotide transport and metabolism | Metabolism | 10 |
| [J] | Translation, ribosomal structure and biogenesis | Information storage and processing | 10 |
| [V] | Defense mechanisms | Cellular processes and signaling | 8 |
| [Q] | Secondary metabolites biosynthesis, transport, and catabolism | Metabolism | 5 |
| [Y] | Nuclear structure | Cellular processes and signaling | 3 |
| [H] | Coenzyme transport and metabolism | Metabolism | 2 |
| [M] | Cell wall/membrane/envelope biogenesis | Cellular processes and signaling | 1 |
| [N] | Cell motility | Cellular processes and signaling | 1 |

**Supplementary Table S10: Pathway enrichment of MSA genes in tiger (XD-score). The top 20 pathway (based on XD-score) are shown after correction.**

| **Annotation** (pathway/process) | **XD-score** | **Fisher q-value** | **Gene set size** | **Pathway size** | **Overlap size** |
|---|---|---|---|---|---|
| **Notch signaling pathway** | **1.4287** | 0.0091 | 846 | 47 | 11 |
| Biosynthesis of unsaturated fatty acids | 0.7579 | 0.7531 | 846 | 21 | 3 |
| Fc gamma R-mediated phagocytosis | 0.6602 | 0.2486 | 846 | 91 | 12 |
| Basal cell carcinoma | 0.6389 | 0.3004 | 846 | 55 | 8 |
| Adherens junction | 0.5988 | 0.3265 | 846 | 72 | 9 |
| Axon guidance | 0.5981 | 0.2221 | 846 | 128 | 16 |
| Other glycan degradation | 0.5972 | 0.9190 | 846 | 16 | 2 |
| Riboflavin metabolism | 0.5972 | 0.9190 | 846 | 16 | 2 |
| Base excision repair | 0.5631 | 0.7531 | 846 | 33 | 4 |
| Insulin signaling pathway | 0.5468 | 0.2367 | 846 | 134 | 16 |
| mTOR signaling pathway | 0.5310 | 0.5524 | 846 | 51 | 6 |
| Endometrial cancer | 0.5310 | 0.3476 | 846 | 52 | 7 |
| Long-term potentiation | 0.5310 | 0.4638 | 846 | 68 | 8 |
| Galactose metabolism | 0.5107 | 0.8794 | 846 | 26 | 3 |
| alpha-Linolenic acid metabolism | 0.4722 | 0.9741 | 846 | 18 | 2 |
| Melanogenesis | 0.4722 | 0.3380 | 846 | 99 | 11 |
| Phototransduction | 0.4722 | 0.8794 | 846 | 27 | 3 |
| Lysosome | 0.4472 | 0.3265 | 846 | 120 | 13 |
| B cell receptor signaling pathway | 0.4452 | 0.5147 | 846 | 74 | 8 |
| Glycosaminoglycan degradation | 0.4196 | 1.0000 | 846 | 19 | 2 |

**Supplementary Table S11: Tissue XD-score based on expression data of genes belonging to Notch pathway.**

| **Tissue type** | **Tissue XD-score** | **Tissue type** | **Tissue XD-score** |
|---|---|---|---|
| **temporal lobe** | 8.62 | amygdala | 1.78 |
| **whole brain** | 8.56 | parietal lobe | 1.69 |

| | | | |
|---|---|---|---|
| **cerebellum peduncles** | 8.21 | subthalamic nucleus | 1.65 |
| **prostate** | 8.08 | skeletal muscle | 1.64 |
| cerebellum | 4.44 | superior cervical ganglion | 1.49 |
| skin | 3.88 | trigeminal ganglion | 1.35 |
| placenta | 2.78 | medulla oblongata | 1.27 |
| globus pallidus | 2.42 | pons | 1.2 |
| atrioventricular node | 2.24 | cingulate cortex | 1.14 |
| liver | 2.22 | dorsal root ganglion | 1.11 |
| ciliary ganglion | 1.85 | prefrontal cortex | 0.79 |
| occipital lobe | 1.8 | testis | 0.47 |

### SUPPLEMENTARY TEXT

#### Supplementary Text S1: Need for error correction

An orthologous gene set of nine species including tiger, constructed using Best reciprocal blast hit (BRBH) approach [1], was analysed to identify the genes showing branch-site positive selection, higher branch dN/dS, high nucleotide divergence (root-to-tip branch length), and unique amino acid substitutions with significant functional impact in tiger. A total of 46 genes showed the above mentioned multiple signs of adaptation in tiger, of which 19 genes belonged to the neuronal development and functioning. Among these, LZTS3 (Leucine Zipper Tumor Suppressor Family Member 3), a gene considered to play a crucial role in regulating the postsynaptic density of synapses, showed multiple signs of adaptation and a phylogenetically corrected correlation between NdN and brain weight (**Supplementary Figure 1**).

The identified unique amino acid substitutions in the 19 neuronal genes were mapped to the corresponding tiger genome assembly (http://tigergenome.org) and were found to be in accordance with the genome assembly. Further, these genes were manually evaluated for the existence of the identified substitutions by mapping genomic reads onto the genome assembly and visualized in Integrated Genome Viewer [2]. In all these 19 genes, we observed that almost all of the substitutions were due to errors in the genome assembly. Furthermore, a similar validation was carried out using the latest version of the tiger assembly and gene set available at Ensembl release 94 (PanTig1.0). In this assembly version also, all the 19 genes showed similar erroneous substitutions. For example, two unique substitutions V and L at positions 362 and 363, respectively, in the LZTS3 gene were a result of errors in the assembly (**Supplementary Figure S1**). In this gene, the codon GAG was replaced with GTG and GTG was replaced with CTG in the (-) strand. Similarly, eight of nine unique substitutions in BEX3 gene were identified as errrors in the assembly (**Supplementary Figure S2**). These observations suggests that the tiger genome assembly reported by Cho et al. 2013 [3] and at Ensembl release 94 comprised of several erroneous bases which were perhaps introduced by the *de novo* assembler or by the read correction tools. These genome assembly errors were primarily single nucleotide changes, of which many were present in the coding regions resulting in synonymous and non-synonymous changes. These errors led to misleading results in our evolutionary analysis performed using this tiger assembly and gene set.

*Methodology*

Nine species from the order Canivora namely, *Mustela putorius*, *Canis familiaris*, *Felis catus*, *Lynx pardinus*, *Panthera tigris*, *Panthera uncia*, *Panthera pardus*, *Panthera leo* and *Panthera onca* were considered to identify evolutionary signatures in tiger. The gene sets for *M. putorius*, *C. familiaris* and *F. catus* were retrieved from Ensembl release 90. The gene orthologs for the five Panthera species were provided by Figueiro et al., 2017 [4]. The gene set for lynx was provided by Abascal et al., 2016 [5]. An orthologous gene set was constructed using BRBH approach [1] considering cat as a reference. The gene phylogeny of each ortholog was inferred from the species phylogeny. The protein alignment of each ortholog was carried out using SATé-II [6], which implemented PRANK for alignment, Muscle for merging the alignment and RAxML for tree estimation. The protein-based nucleotide alignment was carried out using TRANALIGN in EMBOSS package [7]. The variation in ω ratio between lineages on individual genes was calculated using the free-ratio model in CodeML from the PAML software package (v4.9a) [8]. The dN/dS values were calculated for the genes, which qualified likelihood ratio test using a conservative 5% false-discovery-rate criterion against the null model (One ratio). The genes showing higher dN/dS in tiger were identified using the branch model in PAML and the genes which qualified likelihood ratio test against null model (One ratio) were considered to show higher branch dN/dS. For investigation of positive selection, the branch-site model was used in PAML and the likelihood ratio was compared against the null model with 5% false discovery rate. Unique amino acid substitutions in tiger were identified using In-house scripts. Each of the identified substitutions were tested for their functional impact using Sorting Intolerant From Tolerant (SIFT) [9] using UniProt databse as reference. The nucleotide divergence rate was calculated using root-to-tip branch length. The genes showing more than one sign of adaptation were considered for further analysis. The genes evolving with brain size were identified using phylogenetically corrected generalised least square fit in BayesTraits (v3.0.1) [10] (continuous regression) between selection pressure (NdN) and brain size.

**Supplementary Text S2**: **Impact of assembly errors in evolutionary analysis**

Since in our analysis, the incorrect assembly led to several misleading insights in the evolutionary context, we hypothesized that the results from previous studies which used incorrect tiger assembly could also be due to assembly artefacts. Thus, we evaluated the results of such studies in the light of corrected tiger genome assembly.

Figueiro et al., 2017 [4] reported a gene "ESRP1" to be positively selected in jaguar affecting craniofacial robustness. They identified a unique substitution I298Y in jaguar with respect to other species, including tiger (incorrect assembly). We found that the "I" at this position in the tiger gene was an error and is "Y" in the corrected assembly. Moreover, leopard, cat, dog and other mammals also contain "Y" at this position (**Supplementary Figure S3** and **S4**). Further, the study reports six genes from the Glypican pathway to be positively selected in tiger (using branch-site model). Among these six genes, we found that five genes namely, ARNT, GNA15, SIN3B, CA9 and TFDP1 had incorrect bases in their coding sequence, and were corrected in our study. These genes were not found to be positively selected after correction in our study. The authors further identified two genes, DOCK3 and COL4A5 to show the strongest signal of interspecies and intraspecies positive selection in jaguar after performing Bonferroni correction. We found that both these genes had incorrect bases in tiger. These genes were found to show interspecies introgression, which was confirmed by pair-wise divergence assesment in each Panthera species with cat in the windows of 100kb containing these genes. However, we found that a total of 226 bases were erroneous in the selected windows and the rate of error in these windows was 1.5 times greater than the average rate of the error in the complete tiger genome assembly. Thus, these results suggest that substantial corrections have been carried out in the studied region which may significantly affect the interspecies adaptation and introgression analysis for these genes among *Panthera* and needs re-evaluation.

Dobrynin et al., 2015 [11] used the incorrect tiger genome assembly and gene set and calculated a distribution of genome-wide dN/dS values and compared it with *Acinonyx jubatus* (cheetah) genome. The results from this analysis suggest that cheetah distribution is significantly different from other species including tiger. However of these, 4,472 genes have changed in the corrected tiger assembly which will change the distribution of dN/dS across the coding genome. However, we could not verify this due to the inaccessibility of the cheetah genome assembly and gene set.

Montague et al., 2014 [12] carried out evolutionary analysis to identify genes showing adaptive evolution in cat, felids and carnivores. The analysis was performed using the erroneous tiger genome assembly and identified 331 genes to be positively selected in felids, of which 294 genes could be compared with the present study based in their unambiguous assignment to ensembl gene ID. Of these genes, 80 genes had erroneous substitutions in tiger. Our analysis identified only 44 genes to be positively selected (branch-site) or showed higher branch dN/dS (two branch model) in felids using branch-site model among the reported genes (Montague et al., 2014). The study also reports an enrichment of sensory perception related genes among PSGs in felids. Among the 14 genes reported (Montague et al., 2014) to show positive selection in felids with respect to sensory perception, only two genes (GUCA1A and GRIN2C) were found to be positively selected in felids after correction in tiger gene set.

*Effect of asssembly errors on divergence time*

A large number of erroneous bases were identified in this study, which can have significant impact on the divergence time calculation of tiger with its ancestors/other species. Using a mutation rate of 1.1 x $10^{-9}$ per base per year and genome size of 2.4 x $10^{9}$ [3], the mutation rate per year is 2.64, calculated as:

mutation rate per year = mutation rate per base per year x genome size

A total of 982,606 bases were found erroneous in the tiger genome assembly. The upper limit by which these errors can affect the divergence time can be calculated as :

upper limit = erroneous bases x mutation rate per year

Using the above formula, it can affect the divergence time by 0.37 million years.
